# Linking age changes in human cortical microcircuits to impaired brain function and EEG biomarkers

**DOI:** 10.1101/2025.07.03.663090

**Authors:** Alexandre Guet-McCreight, Shreejoy Tripathy, Etienne Sibille, Etay Hay

## Abstract

Human brain aging involves a variety of cellular and synaptic changes, but how these changes affect brain function and signals remains poorly understood due to experimental limitations in humans, meriting the use of detailed computational models. We identified key human cellular and synaptic changes occurring with age from previous studies, including a loss of inhibitory cells, NMDA receptors, and spines. We integrated these changes into our detailed human cortical microcircuit models and simulated activity in middle-age (∼50 yrs) and older (∼70 yrs) microcircuits, and linked the altered mechanisms to reduced spike rates and impaired signal detection. We then simulated EEG potentials arising from the microcircuit activity and found that the emergent power spectral changes due to these aging cellular mechanisms reproduced most of the resting-state EEG biomarkers seen in human aging, including reduced aperiodic offset, exponent, and periodic peak center frequency. Using machine learning, we demonstrated that the changes to the cellular and synaptic aging mechanisms can be estimated accurately from the simulated EEG aging biomarkers. Our results link cellular and synaptic mechanisms of aging with impaired cortical function and physiological biomarkers in clinically-relevant brain signals.

## Introduction

Human aging involves a variety of neurophysiological changes at the cortical microcircuit level, including loss of inhibitory cells^1^, synaptic composition changes^2^, and synaptic spine loss^3^. However, the contributions of these changes to age-related impairments in human brain function, such as executive function and sensory processing^4^, remain poorly understood and result in gaps in diagnosis and treatments. As our ability to study microcircuits in the living human brain is limited, detailed computational models of human brain microcircuits^5^ offer a powerful tool to mechanistically link microcircuit changes to altered brain function and clinically-relevant brain signals such as electroencephalography (EEG) in aging^6^.

A variety of age-related mechanisms have been identified across scales and species, and several key cellular and microcircuit changes have been characterized in detail in the human brain. Transcriptomic studies reported an association between age and differences in cell type proportions^1^, including reductions in somatostatin-(SST) and vasoactive intestinal polypeptide (VIP) expressing interneurons. Human layer 2/3 pyramidal neurons in older individuals also exhibited reduced N-methyl-D-aspartate (NMDA) GluN2A/B protein levels and NMDA-associated synaptic currents^2^. Starting from early adulthood, there was also a steady decline of dendritic spine counts in human pyramidal neurons^3,7^, in particular thin spines and immature spines (filopodia)^8^. Previous studies also suggest inter-dependencies between these mechanisms, whereby filopodia were shown to mainly contain NMDA receptors and lack AMPA receptors^8,9^, boosting SST-mediated inhibition recovered pyramidal neuron spine counts in aged rodents^10^, and long-term dampening of SST-mediated inhibition reduced NMDA-mediated excitation^11^.

These mechanisms have been linked to human and rodent age-related neurological disease and cognitive decline^10,12^. Using novel pharmacology to restore loss of SST cell signaling in aged rodents, which can occur due to increased neuroinflammation^13,14^, hypermethylation^15^ and endoplasmic reticulum stress^16^, was sufficient to recover working memory^10^. NMDA loss has been linked to age-dependent decline of slow inward currents, which modulates spike timing-dependent plasticity with age^17^. In addition, pyramidal neuron spine loss was greater in cognitively impaired older rats compared to unimpaired older and younger rats^18^.

Given the limitations in collecting *in vivo* neuronal and microcircuit data from living humans, non-invasive EEG recordings remain one of the most useful methods for measuring human neuronal activity in health and disease. Several previous studies found consistent and robust age-associated changes in human EEG power spectral features, whereby power spectral decomposition into aperiodic (broadband) and periodic components showed reductions in aperiodic slope and power^6,19^ and reductions in periodic peak alpha frequency and power^6,19–21^. Despite the robust findings about EEG biomarkers in aging, the links to the underlying changes in neuronal and microcircuit mechanisms remain unknown.

Age-associated cognitive decline in older adults includes slower cortical processing^4^, reduced working memory capacity^22^, and reduced discrimination acuity^23,24^. Studies in monkeys have also shown that a decline in working memory with age was associated with reduced baseline and response spike rates in prefrontal cortex^25^. Correspondingly, age-associated declines in spike rates were reported in multi-electrode array data obtained from human cortical slices^26^. However, the link to underlying mechanisms remains unclear.

To overcome experimental limitations in linking altered microcircuit and cellular mechanisms to cognitive impairment and EEG in human aging, we integrated aging mechanisms from human cellular studies into our previous detailed models of human cortical microcircuits. Using these models, we simulated EEG and microcircuit spiking activity to delineate the contributions of cellular aging to microcircuit function, signal integration, and EEG biomarkers of aging.

## Results

We simulated the effect of aging mechanisms on detailed models of prototypical human cortical microcircuits, representing microcircuits of middle-aged (∼50 yrs old) individuals (**Fig. 1A**). The microcircuits included four key neuron types (Pyr neurons and SST, PV and VIP interneurons) constrained with human data of their firing properties, morphologies, cell proportions, synaptic properties and connection probabilities (**Fig. 1A-B**). We integrated cellular and synaptic aging mechanisms from human studies to simulate microcircuits of older individuals. These included loss of inhibitory cells derived from transcriptomic data^1^, loss of Pyr neuron NMDA receptors derived from protein level and synaptic current data^2^, and loss of Pyr neuron spines derived from imaging data^3^, estimated over a 20-year span (i.e., ∼50 yrs vs ∼70 yrs; **Fig. 1B-E**). The experimental changes were modeled linearly as proportional changes in the model’s SST, VIP, and PV inhibitory cell counts (**Fig. 1C**), the NMDA conductance onto postsynaptic Pyr neurons (**Fig. 1D**), and the connection probability and passive parameters of Pyr neurons (reflecting spine loss; **Fig. 1E**).

**Figure 1.**
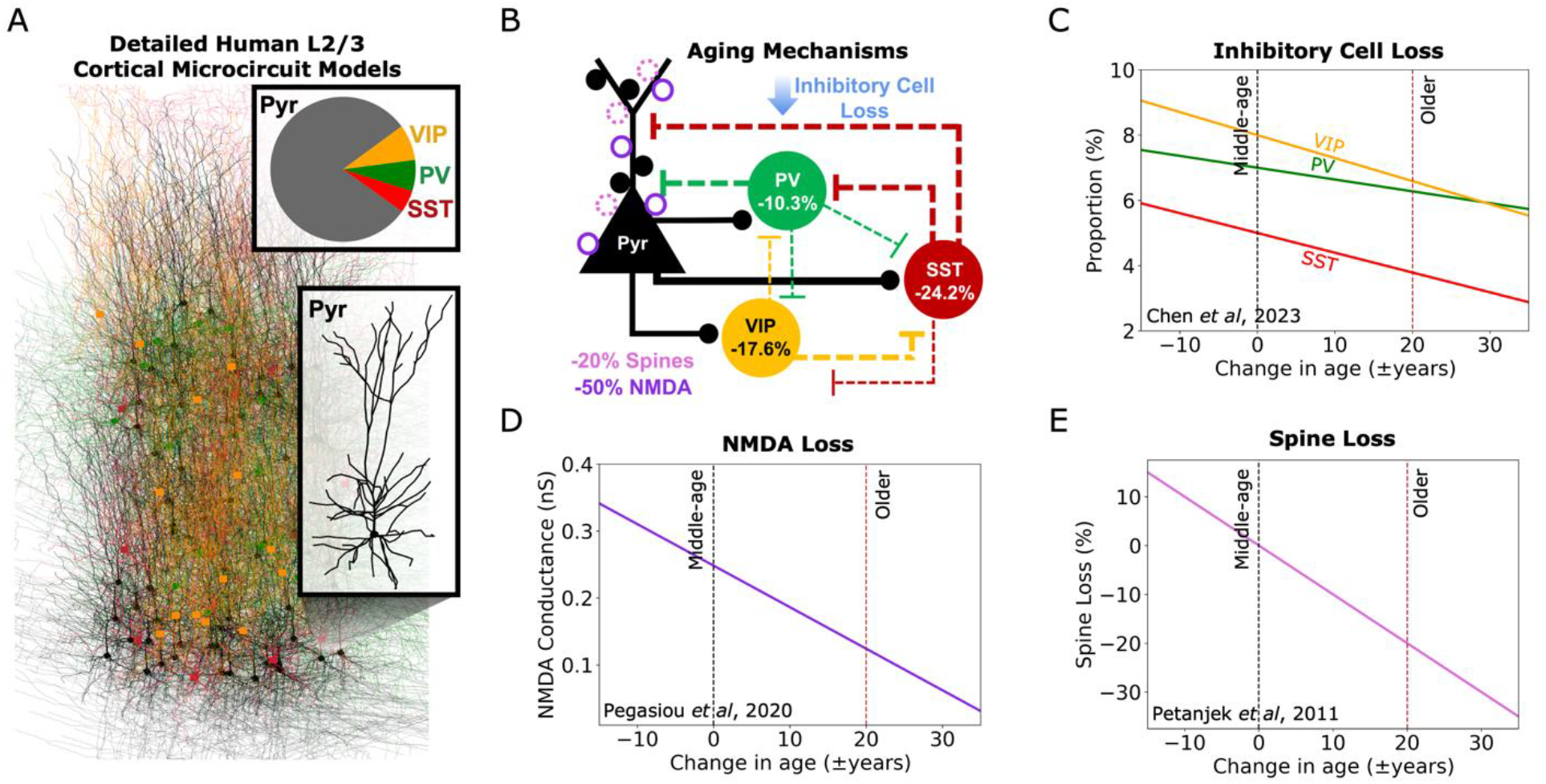
Modeling multiple microcircuit cellular and synaptic mechanisms of human aging. **A**. Illustration of the detailed human L2/3 microcircuit model, which included four key neuron types with human cell proportions, detailed morphologies, firing and synaptic properties. **B**. Schematic of aging microcircuit model, showing connectivity between neuron types and integration of human aging mechanisms data including loss of inhibitory cells, NMDA GluN2A/B protein levels, and spines due to +20 years aging. **C-E**. Data-driven relationship between age and inhibitory cell proportion loss (C), NMDA conductance loss (D) and percent spine loss (E). Dashed lines show middle-age and older (aged +20 yrs) values.

We simulated the middle-aged (∼50 yrs) and older (∼70 yrs) microcircuits, with randomized circuitries to mimic between-person variability, at baseline activity and in response to a brief stimulus (**Fig. 2A**). Older microcircuits had reduced spike rate during baseline (0.76 ± 0.04 Hz vs 0.57 ± 0.03 Hz, *p* = 1.08e-14, Cohen’s *d* = -6.0) and post-stimulus recurrent activity (2.53 ± 0.76 Hz vs 1.74 ± 0.27 Hz, *p* = 8.20e-4, Cohen’s *d* = -1.3; **Fig. 2B**). The dampened activity resulted in higher signal detection errors in older microcircuits, based on readout of recurrent activity vs baseline as measured by the overlap between the spike rate distributions (14.3 ± 2.2% vs 24.9 ± 3.8%, *p* = 1.36e-8, Cohen’s *d* = 3.3; **Fig. 2C-D**). We further simulated activity in response to a stronger stimulus with enhanced recurrent activity and found a larger effect of the aging mechanisms in reducing post-stimulus recurrent activity (7.01 ± 3.49 Hz vs 2.12 ± 0.53 Hz, *p* = 5.64e-6, Cohen’s *d* = -1.9; **Fig. 2E**). With this stimulus an increase in signal detection errors was similarly seen using classifier models (ANN or SVM) to distinguish between baseline and recurrent response activity (**Fig. 2F**). We found that classification accuracy decreased with older microcircuits (ANN: 74.0 ± 20.5% vs 91.2 ± 8.8%, *p* = 1.20e-11, Cohen’s *d* = -1.1; SVM: 71.2 ± 19.1% vs 88.0 ± 10.5%, *p* = 7.40e-11, Cohen’s *d* = -1.1; **Fig. 2G**).

**Figure 2.**
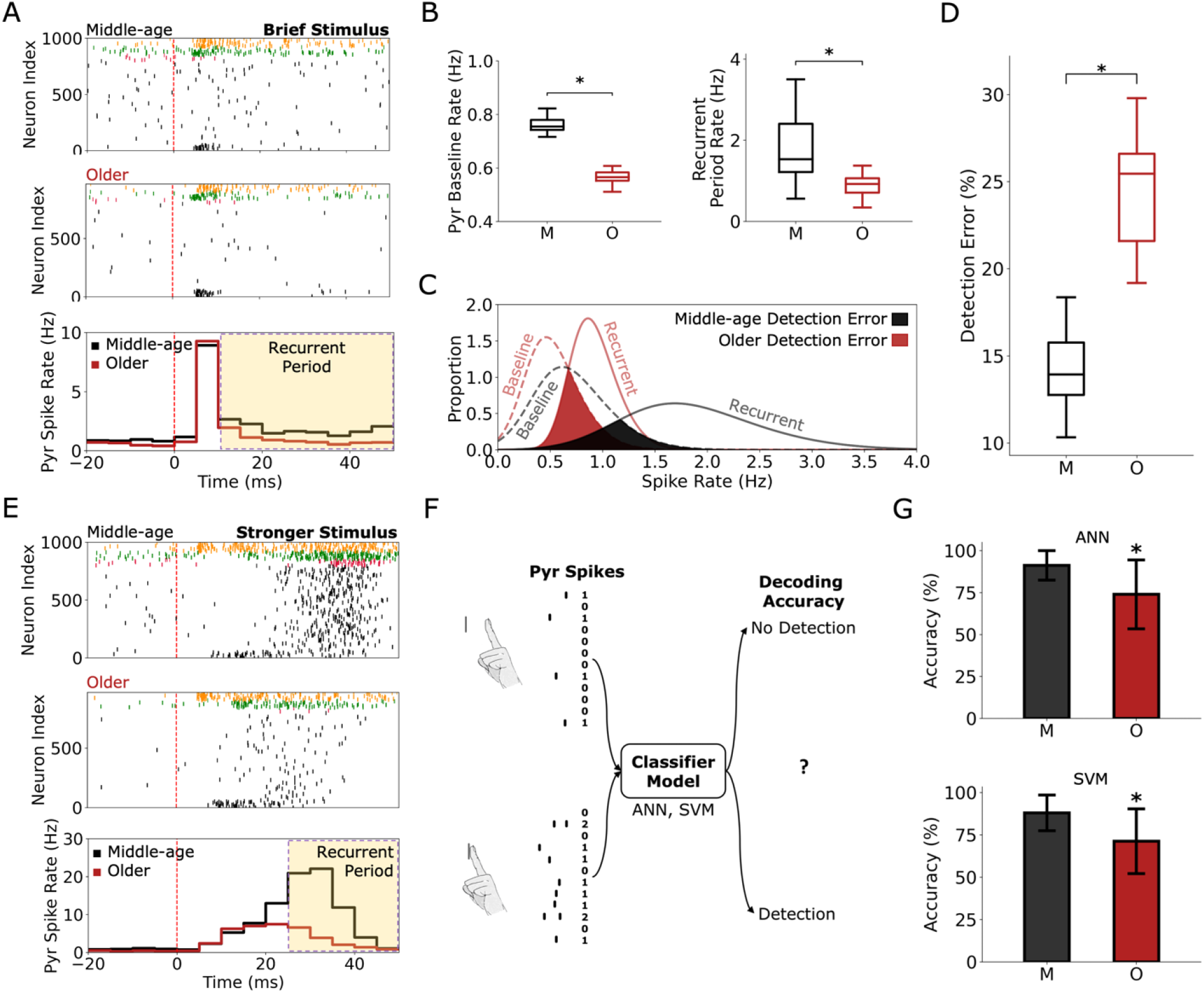
Human aging microcircuit mechanisms reduce spike rates and impair signal processing. **A**. Example raster plots of middle-age and older microcircuits at baseline and in response to a brief stimulus (top), and average peri-stimulus time histograms (PSTH, bottom). Shaded yellow area denotes the time range for the recurrent activity period. Dashed line shows stimulus time. **B**. Mean Pyr neuron spike rates at baseline (left) and recurrent response (right) for middle-age and older microcircuits (20 randomized each). **C**. Distribution of firing rates in the baseline and recurrent response activity periods for the middle-age and older microcircuits. Signal detection error was measured by the percent overlap between the baseline and recurrent period response rate distributions (shaded area). **D**. Signal detection error rate for middle-age and older microcircuits (bootstrapped 1000 permutations). **E**. Example middle-age and older microcircuit raster plots (top) and mean PSTH (bottom) in response to a stronger and de-correlated stimulus. **F**. Schematic of how decoding accuracy was assessed. Spike trains in 50 ms baseline and recurrent period response windows (40 windows total per condition) were summed into spike count vectors (left) and inputted into a classifier model. **G**. Decoding accuracy when using an ANN (top) and a SVM (bottom) for the decoding (n = 100 randomized classifiers). Asterisks denote significant t-tests and Cohen’s *d* > 0.5 of older microcircuits compared to middle-age microcircuits. For box-and-whisker plots, boxes show interquartile range (IQR), middle lines show the medians, and whiskers show 1.5x the IQR.

We next assessed the effect of the aging mechanisms on the simulated EEG from the microcircuit models (**Fig. 3A**), in terms of EEG power spectral density (PSD, **Fig. 3B**), and found a left-shift of peak α frequency (see periodic component analysis below) and a reduced power in θ, α, β frequency bands as measured by AUC (θ: 0.066 ± 0.007 μV^4^/Hz^2^ vs 0.034 ± 0.004 μV^4^/Hz^2^, *p* = 2.17e-12, Cohen’s *d* = -5.7; α: 0.065 ± 0.005 μV^4^/Hz^2^ vs 0.057 ± 0.007 μV^4^/Hz^2^, *p* = 2.50e-4,

**Figure 3.**
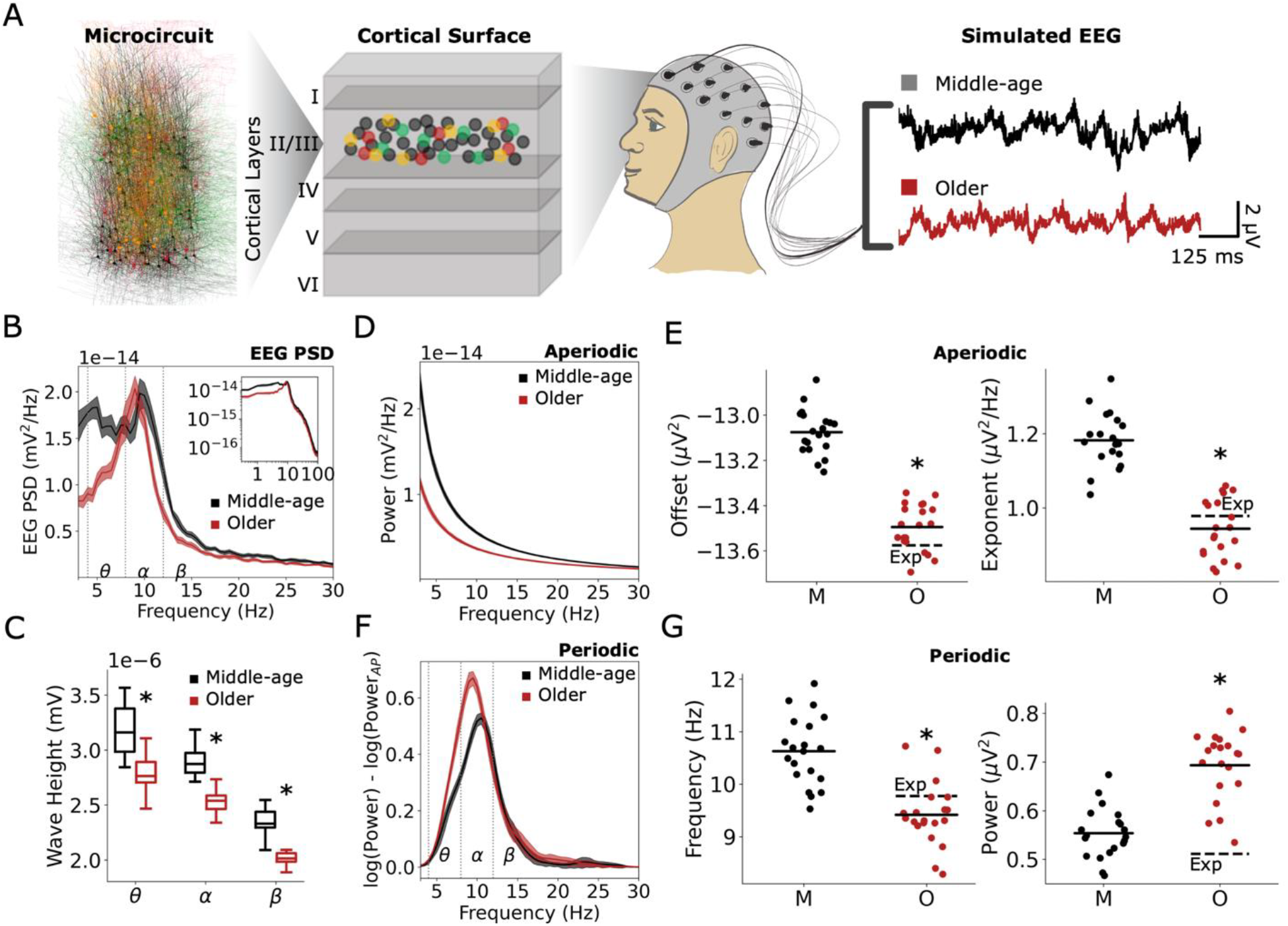
Human aging microcircuit mechanisms account for key EEG biomarkers. **A**. Schematic of simulated EEG signals generated from the human microcircuit models. **B**. Power spectral density (PSD) of simulated EEG from middle-age and older microcircuit models (bootstrapped mean, and 95% confidence intervals). Inset – PSD in log scale. **C**. Wave height extracted from oscillation events and separated by frequency bands. **D-E**. Aperiodic components of the PSD (D) and associated parameters (E, left: offset, right: exponent, dots correspond to n = 20 simulated microcircuits in each age condition). **F-G**. Periodic components of the PSD (F) and associated parameters (G, left: peak centre frequency, right: peak amplitude). In E and G, the dashed lines (Exp) indicate the relative expected changes based on previously reported changes in human aging (Merkin *et al*, 2023). Asterisks and box-and-whisker plots are as described in Fig 2. n = 20 randomized microcircuit simulations per condition.

Cohen’s *d* = -1.3; β: 0.081 ± 0.006 μV^4^/Hz^2^ vs 0.063 ± 0.006 μV^4^/Hz^2^, *p* = 4.11e-10, Cohen’s *d* = -3.0). Correspondingly, in performing a wavelet-based spectrogram event analysis we found decreased wave heights (θ: 3.18 ± 0.22 nV vs 2.79 ± 0.16 nV, *p* = 2.56e-7, Cohen’s *d* = -2.0; α: 2.89 ± 0.12 nV vs 2.53 ± 0.12 nV, *p* = 2.05e-8, Cohen’s *d* = -3.0; β: 2.34 ± 0.12 nV vs 2.03 ± 0.09 nV, *p* = 1.29e-8, Cohen’s *d* = -3.0, **Fig. 3C**).

To better analyze the changes in PSD, we decomposed it to aperiodic and periodic components. The aperiodic component (**Fig. 3D**) in older microcircuits exhibited decreased offset (-13.08 ± 0.10 µV^2^ vs -13.49 ± 0.10 µV^2^, *p* = 1.63e-11, Cohen’s *d* = -4.2, **Fig. 3E**) and exponent (1.18 ± 0.07 µV^2^/Hz vs 0.94 ± 0.08 µV^2^/Hz, *p* =3.42e-9, Cohen’s *d* = -3.1; **Fig. 3E**), consistent with changes seen experimentally^19^. The periodic component of the PSD (**Fig. 3F**) exhibited decreased peak centre frequency (10.63 ± 0.64 Hz vs 9.42 ± 0.59 Hz, *p* = 4.68e-7, Cohen’s *d* = - 1.9, **Fig. 3G**), consistent with changes seen experimentally^19^. However, it also exhibited increased periodic peak amplitudes (0.55 ± 0.05 vs 0.69 ± 0.07, *p* = 1.80e-8, Cohen’s *d* = 2.24; **Fig. 3G**), which was inconsistent with the expected decrease previously reported. Importantly, these changes in EEG emerged simply from implementing the cellular and synaptic aging mechanisms, and were not explicitly optimized or tuned for.

We characterized the effects of individual aging cellular and synaptic mechanisms on microcircuit spiking. While both NMDA loss and spine loss reduced baseline spike rates relative to middle-aged microcircuits (0.47 ± 0.02 Hz, *p* = 2.31e-19, Cohen’s *d* = -10.0; 0.64 ± 0.03 Hz, *p* = 3.70e-10, Cohen’s *d* = -3.5, **Fig. 4A**), inhibitory cell loss increased baseline spike rates (1.02 ± 0.05 Hz, *p* = 1.38e-15, Cohen’s *d* = 5.7). In response to a brief stimulus, both NMDA loss and spine loss decreased recurrent response period rates (1.85 ± 0.41 Hz, *p* = 1.91e-3, Cohen’s *d* = - 1.1; 1.97 ± 0.27 Hz, *p* = 0.005, Cohen’s *d* = -0.9, **Fig. 4B, left**), with no significant effect of inhibitory cell loss, but only spine loss worsened signal detection error (19.76 ± 3.95%, *p* = 1.58e-4, Cohen’s *d* = -1.6; **Fig 4B, right**). In response to the stronger stimulus, only spine loss decreased recurrent response rate relative to middle-aged microcircuits (3.00 ± 0.48 Hz, *p* = 1.01e-4, Cohen’s *d* = -1.6; **Fig 4C**), and none of the mechanisms had an effect of decreasing classification accuracy individually.

**Figure 4.**
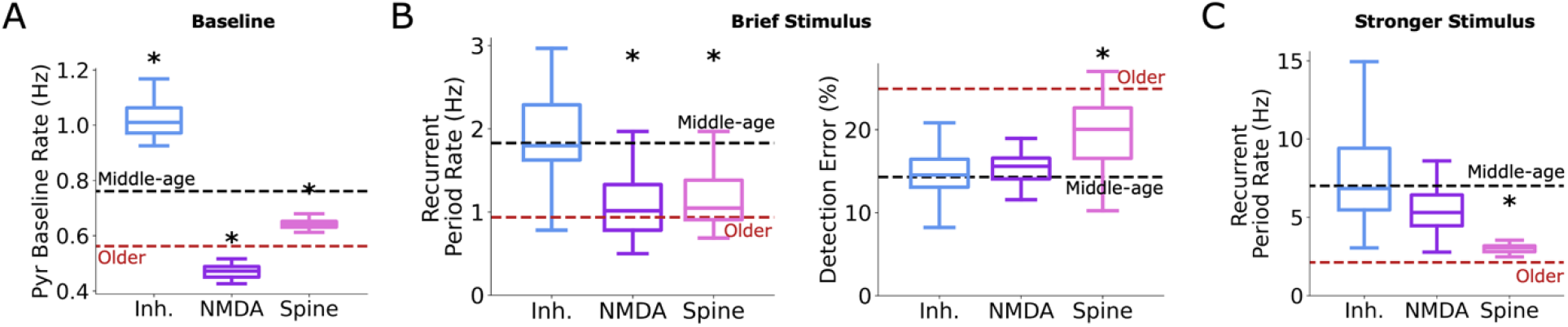
Individual aging cellular and synaptic mechanisms have distinct effects on microcircuit baseline and response activity. **A**. Mean baseline Pyr neuron spike rates for older microcircuits with each aging mechanism implemented separately, compared to middle-age (black dashed lines) and combined mechanisms (red dashed lines). **B**. Recurrent response to a brief stimulus (left) for each aging mechanism and corresponding signal detection errors (right). **C**. Recurrent response to a stronger stimulus for each aging mechanism. n = 20 randomized microcircuit simulations per condition. Box-and-whisker plots are as described in Fig 2.

We next characterized the contributions of individual aging mechanisms to the above EEG biomarkers. All three mechanisms decreased aperiodic offset relative to middle-aged microcircuits (inhibitory cells: -13.08 ± 0.10 µV^2^, *p* = 9.42e-3, Cohen’s *d* = -1.0; NMDA: -13.27 ± 0.09 µV^2^, *p* = 2.59e-6, Cohen’s *d* = -2.1; spine: -13.40 ± 0.10 µV^2^, *p* = 2.51e-9, Cohen’s *d* = -3.2, **Fig. 5A**). However, only inhibitory cell loss and spine loss decreased aperiodic exponents (inhibitory cells: 1.03 ± 0.12 µV^2^/Hz, *p* = 3.01e-4, Cohen’s *d* = -1.5; spine: 0.96 ± 0.08 µV^2^/Hz, *p* = 1.63e-8, Cohen’s *d* = -2.8, **Fig. 5B**). In contrast, only inhibitory cell loss and NMDA loss decreased periodic peak centre frequency (inhibitory cells: 9.60 ± 1.16 Hz, *p* = 1.70e-3, Cohen’s *d* = -1.1; NMDA: 8.92 ± 0.52 Hz, *p* = 5.96e-11, Cohen’s *d* = -2.8, **Fig. 5C**). We additionally characterized the effect on aperiodic area-under-the-curve (1/*f* AUC), for which only NMDA loss and spine loss decreased relative to middle-aged microcircuits (middle-aged: 0.130 ± 0.011 μV^4^/Hz^2^; NMDA: 0.087 ± 0.007 μV^4^/Hz^2^, *p* = 1.17e-12, Cohen’s *d* = -4.6; spine: 0.099 ± 0.007 μV^4^/Hz^2^, *p* = 9.49e-9, Cohen’s *d* = -3.2, **Fig. 5D**). 1/*f* AUC showed a similar direction and magnitude of the aging mechanisms effects as the baseline spike rates (**Fig. 4A** and **Fig. 5D**). In line with this, across conditions we found that 1/*f* AUC correlated strongly with baseline spike rates (Pearson R = 0.85, *p* = 9.59e-30).

**Figure 5.**
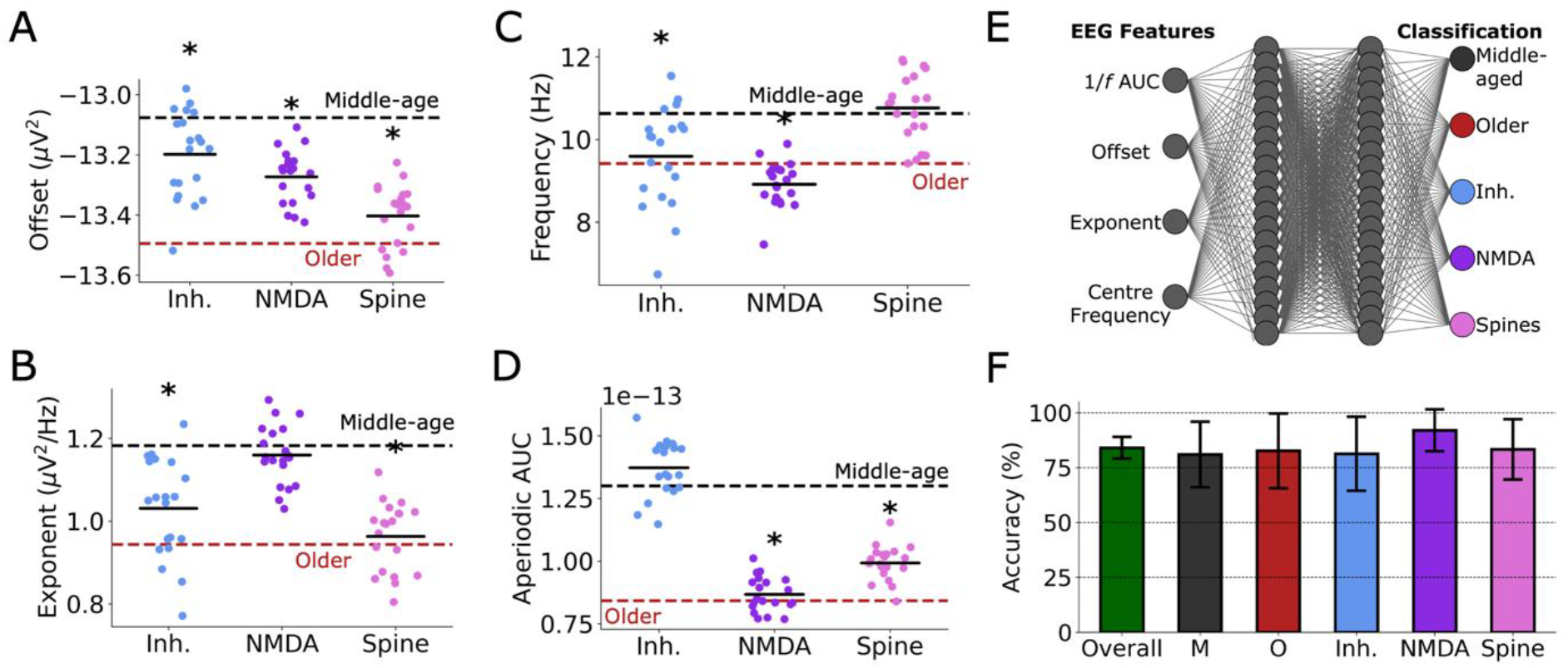
EEG biomarkers of microcircuit aging cellular and synaptic mechanisms. **A-D**. Aperiodic offset (A), exponent (B), periodic peak centre frequency (C), and 1/*f* AUC parameters of each aging mechanism compared to middle-age (black dashed lines) and combined mechanisms (red dashed lines). All asterisks denote significant paired t-tests (p < 0.05) with effect sizes greater than 0.5, when compared to middle-age microcircuits (black asterisks). n = 20 randomized microcircuit simulations per condition. Box-and-whisker plots are as described in Fig 2. **E**. ANN architecture for predicting changes in the cellular and synaptic aging mechanisms from key EEG features. **F**. Test classification accuracies overall and per condition. Accuracy was computed across n = 50 ANN using permutations of 60/10/30% train/validate/test datasets.

Given the differential effects of age-related cellular and synaptic mechanisms on EEG, we assessed whether an artificial neural network (ANN) could classify changes in the cellular and synaptic aging mechanisms based on our simulated EEG biomarkers (**Fig. 5E**). We found high classification accuracy for individual or combined mechanisms in older microcircuits (mean accuracy: 84.1 ± 5.0%, range: 73.3 - 93.3%; **Fig. 5F**). The ANN performed this classification by relying mainly on exponent (SHAP: 0.09 ± 0.02), 1/*f* AUC (SHAP: 0.09 ± 0.02), and offset (SHAP: 0.08 ± 0.03), with a smaller contribution from centre frequency (SHAP: 0.05 ± 0.02).

## Discussion

In this work, we integrated aging mechanisms into detailed models of human cortical microcircuits, and showed that key cellular and circuit human aging mechanisms can account for age-associated changes in spiking and EEG biomarkers. Importantly, the EEG effects emerged from simply implementing the cellular and synaptic aging mechanisms, and were not explicitly tuned for. We further demonstrated that the cellular and synaptic aging changes could be estimated from simulated EEG power spectral biomarkers using artificial neural networks. Our findings overcome challenges in linking cellular and synaptic aging mechanisms with impaired cortical function and aging biomarkers in brain signals, which can enable a more mechanistic stratification of older individuals with age-associated cognitive decline.

The simulated aging EEG effects that we found reproduce most of the biomarkers that have been observed and replicated in several studies, including reduced aperiodic exponent, offset, and periodic peak center frequency^6,19^. Our results are also in line with previous work that showed accurate estimation of age from EEG^27,28^. Here we provide the first link between the EEG effects and human cellular and synaptic aging mechanisms and show that each mechanism generated unique effects. In line with reduced aperiodic parameters due to inhibitory cell loss, boosting inhibition with either pentobarbital or diazepam in rodents has been shown to increase aperiodic exponent and offset^29^. This was also seen in macaques during propofol administration, and was predicted by computational simulations of reduced E/I balance via increased inhibition^30^. Changes in aperiodic exponents with age have also been linked to cognitive decline^21^, a relationship that is influenced by level of education^31^. In line with our effects of inhibitory cell and NMDA loss on alpha peak center frequency, blocking inhibition with picrotoxin or blocking NMDA with ketamine both lead to reduced peak periodic frequencies in the 6-10 Hz range in rodents^29^. Decreased alpha peak frequencies in aging have also been associated with working memory deficits^32^. While no studies to date have directly characterized the effects of dendritic spine changes on EEG, spine density is decreased in rodent medial prefrontal cortex during corticosterone administration^33^, which has also been associated with decreased aperiodic exponent and offset^34^, in line with the effects of spine loss in our models. Interestingly, the alignment in the direction of effects across mechanisms between 1/*f* AUC and baseline spike rate that we found suggests that the dynamics of both measures is similarly sensitive to changes in these underlying neurophysiological mechanisms. In addition, the strong correlation between 1/*f* AUC and baseline spike rate indicates that the simulated resting state activity involves mostly aperiodic, noisy, spiking.

The aging mechanisms led to reduced baseline and response spike rates, in line with effects reported in human cortical slices^26^ and *in vivo* in monkeys during a working memory task^25^. These spiking changes resulted in worsened signal detection errors, in agreement with impaired cognitive performance during sensory discrimination in older subjects^24^. Baseline spike rates were increased by inhibitory neuron loss, consistent with our previous work on the effects of SST inhibition loss^5^, and decreased by NMDA and spine loss, as expected from reduced excitation onto Pyr neurons^35,36^. While inhibitory neuron loss had no effect on response rates (consistent with our previous work^5^), both NMDA and spine loss reduced recurrent responses to brief stimuli, consistent with previous findings on excitatory effects during brief thalamocortical stimulation^36^ and ketamine application *in vivo*^37^. Conversely, spine loss but not NMDA loss reduced recurrent responses to stronger stimuli, possibly due to dendritic saturation effects (i.e., reduced driving force) with larger stimuli^35,38^ causing smaller differences in NMDA output at higher activation levels. We did not disentangle the effects of SST, PV, and VIP interneuron loss, but the EEG changes we report due to inhibitory interneuron loss match more distinctly with EEG changes seen during SST inhibition loss rather than PV inhibition loss^39^, which is in line with the greater magnitude of SST loss that we simulate in our aging models.

While the changes in aging mechanisms that we modeled accounted for several key features of aging EEG, none of the mechanisms alone or together replicated the decrease in periodic peak power seen previously^6,19^, suggesting the involvement of other aging mechanisms. Changes in ion channel balances such as those we reported previously in human L5 Pyr neurons^26^ may contribute to a decrease in periodic peak power. Age-related EEG changes may also be affected by non-neuronal mechanisms, such increased glial cell activity^1^, changes in cardiac activity^40^, genetic risk factors for dementia^21^, or other cardiovascular factors^21^. In addition, for our aging mechanisms we assumed linear reductions with age, but these relationships may be non-linear and future experimental studies will be important for characterizing these relationships more clearly. The changes in periodic peak power with age may also involve components not currently included in the model, such as layer 5, which contributes to beta rhythm generation^26,41^, or active dendritic properties^42,43^. Given the low incidence of beta events in our layer 2/3 models, these were not well suited to capture changes in beta frequency oscillations, which increase in peak power and decrease in peak centre frequency with age^44^. While we simulated a generic prototypical human cortical microcircuit and studied general stimulus response (which can apply to either sensory or higher order regions), future studies may benefit from modeling a specific area of cortex, such as sensory or prefrontal cortex^45^ to study particular age-related impairments. In addition, exploring the effects of cellular and synaptic aging over more time points could provide additional insights into how these mechanisms impact EEG and microcircuit function over time.

While we tested the accuracy of ANN trained for mechanistic classification, another approach would be to train ANN for estimating the level of cellular aging, which would require more biophysical simulations of different severity levels for each mechanism. Although the ANN we used had good classification accuracy using two hidden layers, future studies could conduct a more comprehensive search through network architecture and feature sets to further improve the accuracy or establish the utility of features. It will be of particular interest for future studies to apply our in silico-trained artificial neural networks to estimate cellular aging for human subjects from their EEG data and assess the correspondence with cognitive impairments. Such effort will enable a mechanistic stratification, improved diagnosis, and establish target treatment mechanism in aging.

## Methods

### Models of human cortical microcircuits

We used our previous morphologically- and biophysically-detailed models of human L2/3 cortical microcircuits^5^. The models were comprised of 1000 neurons belonging to key neuron types (80% Pyr, 5% SST, 7% PV, and 8% VIP as estimated from RNA-seq data^46^) distributed across a 500×500×950 µm^3^ volume. Neuron firing and synaptic connection properties were fitted to human data, including Pyr → Pyr^47^, SST → Pyr apical dendrites^48^ and PV → Pyr basal dendrites^49^. Synapses were modelled using presynaptic short-term plasticity parameters for vesicle-usage, facilitation, and depression, and separate rise and decay parameters for the AMPA and NMDA components of excitatory synapses (τ_rise,NMDA_ = 2 ms; τ_decay,NMDA_ = 65 ms; τ_rise,AMPA_ = 0.3 ms; τ_decay,AMPA_ = 3 ms; τ_rise,GABA_ = 1 ms; τ_decay,GABA_ = 10 ms)^36,50,51^. For complete list of data provenance in our models, please refer to our previous work^5^. Simulations were run using NEURON 7.7^52^ and LFPy 2.0.2 (Python 3.7.6)^53^ on SciNet parallel computing^54^. Our microcircuit model represented a prototypical cortical microcircuit of a middle-aged individual (∼50 years), given the diverse human data (morphology, proportions, electrophysiology, and synaptic connectivity) with which the model was constrained involved ages spanning from 18 to 75 years overall^46,47,55,56^, and cell proportions from human data with a median age of 45 years^46^. We compute all age-related mechanistic changes in terms of relative years, including reduced inhibitory cells^1^, reduced NMDA conductance^2^, and reduced spines^3^. In all cases, we based our model aging mechanisms on human data changes over +20 years (i.e., ∼70-year microcircuits).

### Simulating age-dependent inhibitory interneuron loss

For reduced inhibitory cells with age, we analyzed cell type proportions starting from human single-cell RNA-seq data (MSSM and McLean cohorts, age range: 24 – 74 years, n = 71, 30 females)^1^ to extract age regression coefficients and intercepts for each interneuron type in our model (SST, VIP, PV). From these we derived a loss of 1.21% SST interneurons, 0.88% VIP interneurons, and 0.52% PV interneurons per year, and thus 24.2%, 17.6% and 10.3% loss of each interneuron type respectively over 20 years aging from middle-age to older microcircuits.

### Simulating age-dependent NMDA loss

For reduced NMDA conductance with age, we estimated a 2.5% decline per year in human L2/3 Pyr neuron GluN2A/B protein levels from western blot data (age range: 21 – 71 years, n = 17, 9 females)^2^, and thus a 50% decline for aging by 20 years. We translated this as a 50% loss of NMDA conductance in our microcircuit models.

### Simulating age-dependent spine loss

We estimated a 1% spine loss per year in human Pyr neurons from rapid Golgi staining data (age ranges included: 20 – 91 years; n = 18, 4 females)^3^, starting from adulthood (∼20 years). We translated this as a 20% loss of Pyr → Pyr connection probability over 20 years aging between the middle-age and older microcircuits. Additionally, we modeled the resulting loss of dendritic surface area, by reduced dendritic passive conductance (*G*_*pas*_) and membrane capacitance (*c*_*m*_). In humans, spine loss with age is mostly due to a loss of thin spines, which, in older individuals comprise approximately 25% of spines^8^. Given a 1% loss over 20 years, we thus estimated that thin spines in middle-aged individuals comprise approximately 31.25% of spines. Given a total number of 8,169 spines in young monkey Pyr neurons^57^, and assuming a thin spine length of 1.2 μm^58^, a tip diameter of 0.35 μm^58^, and a truncated cone shape with a half base-to-tip diameter ratio, we computed the total surface area due to thin spines to be approximately 2,782.57 μm^2^. Given a total spine surface area of 11,954 μm^2^ and a total dendritic surface area of 20,410 μm^2 57^, we computed the proportion of surface area due to thin spines in middle-age to be 8.6% of the total dendritic and spine surface area. This yielded a reduction of 1.72% in dendritic (basal + apical) *G*_*pas*_ and *c*_*m*_ over 20 years aging between middle-age and older microcircuits.

### Microcircuit baseline and response spiking activity

We simulated baseline microcircuit activity as described previously^59^, where we provided background input to the microcircuit with excitatory Ornstein Uhlenbeck (OU) point processes^60^. Independent excitatory OU point processes were placed midway along the length of each dendritic arbor, and for Pyr neuron models, we placed 5 additional OU processes at 10%, 30%, 50%, 70%, 90% of the apical length. We scaled up the mean and standard deviation of each OU conductance exponentially with relative distance from soma (ranging from 0 to 1) to normalize their effect.

For the brief stimulus, we used previous models of response rates^5^, using excitatory AMPA/NMDA synapses with the same synaptic dynamics as the cortical excitatory synapses. We stimulated the basal dendrites of 55 Pyr neurons, with 2–4 ms delay post-stimulus and a conductance of 4 nS. We also stimulated 35 PV interneurons with a delay of 2–2.5 ms and a conductance of 2 nS. VIP interneurons were stimulated in two groups and phases: early (65 VIP interneurons, delay = 2–2.5 ms, conductance = 2.8 nS) and late (80 VIP interneurons, delay = 7– 12 ms, conductance = 2.2 nS).

For the stronger stimulus, we modified the brief stimulus in several ways. We first removed stimulation to PV interneurons to reduce feedforward inhibition. We also increased disinhibition by increasing the stimulus *G*_*max*_ for early VIP interneurons from 2.8 nS to 5.6 nS and late VIP interneurons from 2.2 nS to 4.4 nS. Lastly, we prolonged excitatory stimulation by increasing the Pyr neuron stimulus delay range from 2–4 ms to 2–22 ms, doubled the number of excitatory synapses to Pyr neurons, and compensated by reducing the *G*_*max*_ for Pyr neurons from 4 nS to 2 nS. The stronger stimulus was thus a combination of reduced feedforward inhibition, increased disinhibition, and prolonged excitatory stimulation.

### Signal detection metrics and decoding accuracy

Similarly to previous work where we computed probabilities of failed and false detections^5^, we calculated signal detection errors as the percent overlap in the distributions of Pyr neuron firing rates at baseline (computed using a 40 ms sliding window, sliding in 1 ms intervals, over a 3 s pre-stimulus period) and the distribution of firing rates during the recurrent activation period in response to the brief stimulus (calculated across 20 stimulus presentation, in the 10-50 ms period post-stimulus).

To assess decoding accuracy of recurrently activated Pyr neurons in response to a stronger stimulus, we summed the spikes of each Pyr neuron during the baseline period (50 ms pre-stimulus) and the recurrent activation period (25-75 ms period post-stimulus). The summed spike trains during baseline or response periods of each Pyr neuron were then used as input features into classifier models available in the scikit-learn Python module. The classifier models were trained on 70% of the pre-/post-stimulus windows to distinguish baseline (noise) from recurrent response (signal), and were then tested on the remaining 30% of the data. To compute accuracy statistics, we used 100 permutations of train/test subsets.

### Simulated microcircuit EEG and power spectral analysis

As in previous work^39^, we simulated resting-state EEG from our microcircuit models in LFPy 2.0.2 (Python 3.7.6) using a four-sphere volume conductor model (representing grey matter, cerebrospinal fluid, skull, and scalp with radii of 79 mm, 80 mm, 85 mm, and 90 mm, respectively). The conductivity for each sphere was 0.047 S m^−1^, 1.71 S m^−1^, 0.02 S m^−1^, and 0.41 S m^−1^, respectively^39,61^. We computed power spectral density (PSD) using Welch’s method^62^ from the Scipy python module with 2s time windows. We also decomposed the EEG power spectra (in the 3–30 Hz range) into periodic and aperiodic components using the FOOOF toolbox^6^. The aperiodic component was a 1/f function parameterized by vertical offset and exponent parameters. We fitted the periodic oscillatory component with up to 3 Gaussian peaks defined by center frequency, bandwidth (min: 2 Hz, max: 6 Hz), and power magnitude (relative peak threshold: 2, minimum peak height: 0)^39,59^. We then extracted exponent and offset parameters from the 1/*f* aperiodic component and maximum peak centre frequency and amplitude from the periodic component. For comparison to literature values in figure 3, we extracted corresponding average FOOOF values from Merkin *et al*, 2023^19^, which compared EEG recordings across the scalp (excluding FP1, FPz, FP2, AF7, and AF8) between younger (n = 85; 18–35 years; 37 male) and older adults (n = 92; 50–86 years; 53 male). From these we computed the expected change in FOOOF metrics relative to the mean FOOOF values of our middle-age microcircuit models. We also performed a wavelet-based spectrogram event analysis using the toolbox OEvents^63^ in python, as described previously^59^ (median threshold = 4.0, sampling rate = 40,000 Hz, window size = 24 seconds, minimum frequency = 1 Hz, maximum frequency = 100 Hz, frequency step = 0.5 Hz, overlapping bounding box threshold = 0.5).

### Machine learning models for classifying changes in cellular and synaptic aging mechanisms from EEG features

We used Tensorflow^64^ to train ANN models to classify age-related losses in inhibitory cell type proportions, NMDA receptors, and spines. The ANN models comprised of 4 input nodes, 2 × 20 hidden layer nodes with ReLU activation, and 5 output nodes with softmax activation (1 for each condition, see below). Input features included offset, exponent, peak centre frequency, and 1/*f* AUC. The model was trained to classify middle-age, older, inhibitory loss, NMDA loss, and spine loss microcircuits. The Adam optimizer with Nesterov momentum and a learning rate of 0.001 was used for learning, and loss was measured using categorical cross-entropy. Weights were initialized from a normal distribution centred at zero. All ANN models were trained using 60% of the data, validated using 10% of the data, and tested using 30% of the data. To enable a robust estimation of changes in the cellular and synaptic aging mechanisms and to assess ANN performance, we trained 50 ANN models using random permutations of the data split into 60/10/30 training/validation/testing. ANNs were trained for 100 epochs and we enabled early stopping if validation loss did not reduce for 10 epochs. SHAP values were used to identify feature importance, where larger magnitudes indicated larger impact on model performance^65^.

### Statistics

For group comparisons we used two-sided paired- and independent-sample t-tests assuming equal variance, where indicated. Cohen’s *d* was calculated as follows:

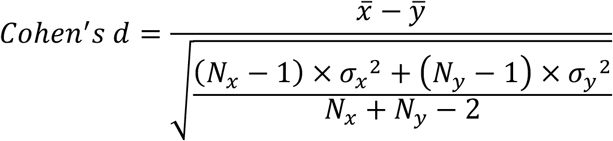

## Code availability

All original code will be deposited on Github as of the date of publication.

## Acknowledgements

AGM and EH thank the Krembil Foundation for their generous funding support. AGM thanks the Canadian Institutes of Health Research - Institute of Aging for funding support.

